# Into the range: a latitudinal gradient or a center-margins differentiation of ecological strategies in *Arabidopsis thaliana*?

**DOI:** 10.1101/2021.10.15.461205

**Authors:** Aurélien Estarague, François Vasseur, Kevin Sartori, Cristina Bastias, Denis Cornet, Lauriane Rouan, Gregory Beurier, Moises Exposito-Alonso, Stéphane Herbette, Justine Bresson, Denis Vile, Cyrille Violle

## Abstract

**Background and Aims:** Determining within-species large-scale variation in phenotypic traits is central to elucidate the drivers of species’ ranges. Intraspecific comparisons offer the opportunity to understand how trade-offs and biogeographical history constrain adaptation to contrasted environmental conditions. Here we test whether functional traits, ecological strategies and phenotypic plasticity in response to abiotic stress vary along a latitudinal or a center-margins gradient within the native range of *Arabidopsis thaliana*.

**Methods:** The phenotypic outcomes of plant adaptation at the center and margins of its geographic range were experimentally examined in 30 accessions from southern, central and northern Europe. The variation of traits related to stress tolerance, resource use, colonization ability as well as survival and fecundity was determined in response to high temperature (34°C) or frost (- 6°C), in combination with response to water deficit.

**Key Results:** Both evidence for a latitudinal and a center-margins differentiation was found. Traits related to the acquisitive/conservative strategy trade-off varied along a latitudinal gradient. Northern accessions presented a greater survival to stress than central and southern accessions. Traits related to a colonization-competition trade-off followed a center-margin differentiation. Central accessions presented a higher phenotypic plasticity and trait values associated with a higher colonization ability than northern and southern accessions which instead had a higher competition ability.

**Conclusions:** Intraspecific phenotypic variation helps us understand how the distribution range has evolved in *Arabidopsis thaliana*, which is shaped both by climate and the population migratory history. We advocate to consider intraspecific trait variation in species range studies instead of species means only as classically done in macroecology.

## Introduction

The way species deploy various ecological strategies to cope with local abiotic and biotic conditions across their geographical distribution range is critical to understand the evolution of distribution range (Brown, 1984; Banta et al., 2012; Schurr et al., 2012). Surprisingly though, most theoretical developments on the determinants of species’ ranges have focused on biogeographical and evolutionary aspects linked to colonization ability (e.g., Kirkpatrick and Barton, 1997; Sexton et al., 2009; Bridle et al., 2010) while overlooking the divergence of populations in term of eco-physiological traits. Phenotypic adaptations along environmental gradients have been widely recognized both between- and within species in functional and evolutionary ecology (Reich et al., 1997, 2003; Wright et al., 2017; Dong et al., 2020; Kuppler et al., 2020). However, such phenotypic adaptations have been scarcely added to the long list of *usual suspects* that determine species’ range size and dynamics (Brown and Gibson, 1983; Gaston, 2009; Sexton et al., 2009). The ecological drivers of range size variation remain largely tackled through the comparison of multiple species while considering species’ ecological characteristics as fixed. This is most often implicit in model species distributions studies (Guisan and Thuiller, 2005), and explicit in studies dedicated to the analysis of phenotypic diversity (namely functional diversity) across species and scales (Violle et al., 2014). The lack of consideration of within-species ecological variation might be necessary in biogeography from a pragmatic point of view, but it ignores theoretical expectations of major differences in ecological performances – survival, growth and reproduction – when moving from the center to the margins of the range of a given species (Abeli et al., 2014; Csergő et al., 2017; Salguero-Gómez et al., 2018).

Plant trait-based ecology has long investigated the variability of phenotypic features (functional traits hereafter) among species, and has linked it to the environments and communities they live in (Violle et al., 2007; Garnier et al., 2016). The joint analysis of the variability of multiple traits further led to the identification of plant ecological strategies that are expected to reflect the phenotypic outcome of natural selection at a given place (Westoby et al., 2002). Notably, based on the combination of a limited number of plant functional traits, the CSR scheme (Grime, 1977, 1988; Hodgson et al., 1999; Pierce et al., 2013, 2017) depicts alternative ecological strategies displayed by any plant species within a triangle whose three summits represent plants completely invested in either competitive strategies (C), stress-tolerant strategies (S), or ruderal strategies (R). Despite its simplicity, the CSR scheme has successfully been used to describe plant community gradients across broad environmental clines (Cerabolini et al., 2010; Rosenfield et al., 2019). However, this classification remains silent regarding its underlying adaptive causes, although this was a prerequisite of plant functional ecology at its infancy (Calow, 1987). The lack of consideration of the adaptive value of functional strategies is partly due to a negligence of intraspecific trait variation in functional ecology (Albert et al., 2010, 2011, 2012; Violle et al., 2012). Recent efforts have emphasized noticeable variations of plant functional strategies across ecotypes of a given species, and demonstrated their adaptive value (Vasseur, et al., 2018a). The exploration of functional trait variation across species’ range is promising since they can reveal the ability of populations to adapt to local, potentially stressful, conditions through functional specialization.

It is expected by definition that the populations at the edges of a species distribution experience the most extreme environmental conditions the species can tolerate (Brown, 1984; Holt, 2009). The contrasted environmental conditions throughout the species’ distribution area are expected to select for differential values of functional traits that reflect physiological tolerance and plant performance as a whole, but also for different levels of phenotypic plasticity. Theoretical considerations predict higher adaptive phenotypic plasticity at the margins of the distribution than at the center (Chevin and Lande, 2011) as a flexible adaptive response to stressful conditions (Chevin and Lande, 2009). Strikingly, the few empirical studies that explicitly quantify plastic divergence across the range draw divergent conclusions, depending on the trait and on the species. In some cases, phenotypic plasticity was found to be higher at the margins than at the center, which was interpreted as an adaptation to more stressful and fluctuating climatic conditions (Volis et al., 1998, 2001, 2015; Lázaro-Nogal et al., 2015; Carvajal et al., 2017). Conversely, some studies highlighted lower plasticity at the margins of plant species’ distribution compared to the center (eg., Mägi et al., 2011), which was explained by a higher cost of maintaining environment sensors in stressful conditions (van Kleunen and Fischer, 2005). Testing hypotheses linking variation and plasticity of functional traits with geography will thus be key to understanding the emergence of species distribution ranges.

The model species *Arabidopsis thaliana* (L.) Heynh., for which both functional traits (Lasky et al., 2012; Vile et al., 2012; Vasseur et al., 2018ab; Sartori et al., 2019; Exposito-Alonso, 2020) as well as biogeographic history (Lee et al., 2017; Hsu et al., 2019) are well studied, presents a unique opportunity to test above hypotheses (Takou et al., 2019). The native distribution of this annual selfing species extends from north Africa to the north of Norway and thus its populations experience dramatically-different environmental conditions (Hoffmann, 2002). Thanks to an international effort of sampling, seeds from more than a thousand of fully sequenced accessions are available (1001 Genomes Consortium, 2016). Taking advantage of this unique genomic database, Lee *et al*. (2017) reconstructed the recent history of colonization of Europe of *A. thaliana*. They showed that the majority of actual european lineages originate from the recolonization of a single lineage of Europe, from central Europe to the south and to the north since the last glacial event. This central lineage then admixed with southern and northern populations. Thus, comparing northern and southern margins allows to compare adaptations to very contrasted climates in a similar demographic context (Lee et al., 2017; Hsu et al., 2019). Moreover, single-nucleotide polymorphisms (SNP) analyses suggested that recolonization of Europe may correlate with adaptations to the contrasted European climates (Méndez-Vigo et al., 2011; Lasky et al., 2012). A huge temperature gradient could be an intense selective strength throughout the latitudinal distribution of *A. thaliana* (Kaplan et al., 2004; Swindell et al., 2007; Vile et al., 2012). Furthermore, some similarities in water availability (due to summer drought or winter frost) may suggest similar strategies for water use on the two opposite margins (Exposito-Alonso et al., 2018). Interestingly, genomic and functional ecology studies provided contrasted evidence in *A. thaliana*. On the one hand, alleles conferring resistance to extreme drought are maintained at both geographical margins (Exposito-Alonso et al., 2018). On the other hand, the “S” (stress-resistance) strategy seem to be displayed by northern accessions only (Vasseur et al., 2018a). Again, the lack of consideration for phenotypic plasticity in functional ecology, and its role in local adaptation, impedes a comprehensive understanding of variation in plant ecological strategies throughout a species distribution range. Here we asked: (i) How do functional traits and strategies vary across the distribution range of *A. thaliana*? (ii) Do the plasticity of traits and strategies differ between the center and the margins of its distribution range? (iii) Are the accessions from the margins more resistant to abiotic stresses than those from the center? To answer these questions, we analyzed functional traits and performance variations of 30 accessions from the south, the center and the north of Europe, grown in controlled conditions under different temperature and water availability treatments.

## Material and methods

### Plant material

We chose thirty natural accessions of *Arabidopsis thaliana*, randomly selected among three geographical groups (**Fig. 1, [Supplementary Information Table S1])**. Ten accessions came from Iberian Peninsula and from Cape Verde, ten accessions from central Europe and ten accessions from Scandinavia (namely hereafter South, Center and North, respectively). All the seeds originated from multiplication realized at the Center of Evolutionary and Functional Ecology (CEFE, Montpellier, France) from original stocks of the 1001Genome Project (1001 Genomes Consortium, 2016). These accessions covered a large range of climatic conditions where *A. thaliana* can grow **[Supplementary Information Fig. S1]**. This set of thirty accessions represents 86.4% of allelic diversity of *A. thaliana*.

**Figure 1.**
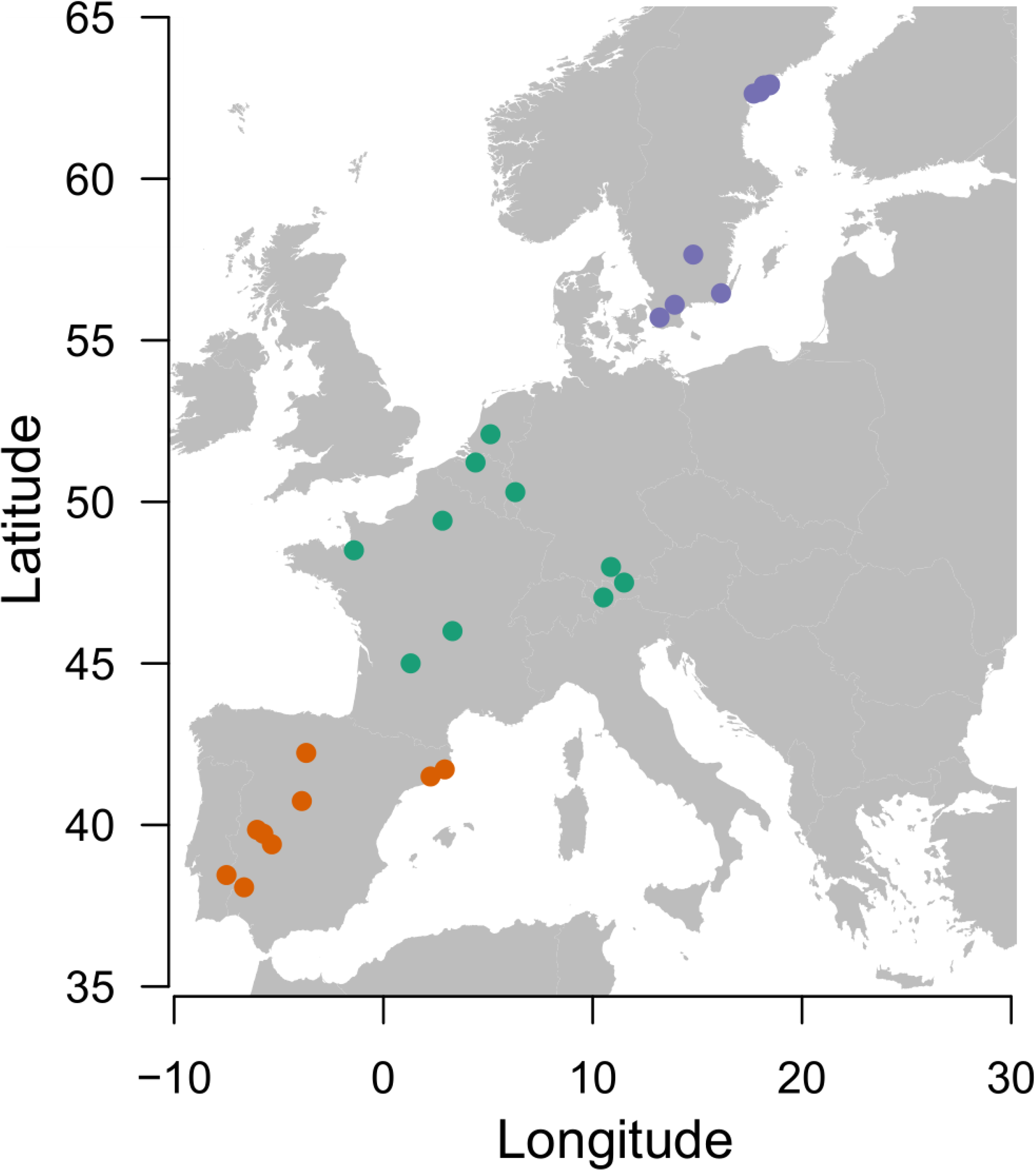
Geographical origin of the 30 accessions of *Arabidopsis thaliana*. The southern group is represented with orange dots, the central group in green and the northern group in purple. Accessions were chosen randomly in each geographical group.

### Experimental design

Seeds of the 30 accessions were sown in November 2018. We used 25 alveolate culture plates containing 120 individual pots of 130mL each filled with peat soil (Neuhaus Humin substrat N2). Each accession was replicated four times in every plate and distributed randomly within and among plates (n = 100 replicates per accession). We stratified seeds by placing plates at 4 °C during for four days. Then, the plates were placed in a greenhouse at 10 °C average temperature during 40 days for vernalization. During vernalization period, we irrigated the pots by osmosed water dipping for 30 minutes once a week. Thereafter, we settled the temperature at 15 °C until the end of January 2019. We then applied five environmental conditions during two weeks (**Table 1)**. These five environmental conditions were composed of five culture plates each, for a total of 20 individuals per accession and per condition (n = 600 individuals in each condition). The control condition consisted of a temperature of 15 °C day and night without any water limitation. These conditions are considered as non-stressful for *A. thaliana*. The cold (LT) treatment consisted of a nocturnal temperature of −6 °C and 15 °C during the day. We set up the nocturnal temperature in a refrigeration enclosure where temperature was homogenous inside (Platinium PLAT7BT, Franstal, France). The Hot (HT) treatment consisted of a daily temperature of 35 °C and of 15 °C during the night. For this treatment, we moved the plants in another compartment at 35 °C. Light and air humidity were kept identical both in LT and HT treatments. In each temperature condition, half of the plates was sub-irrigated at field capacity once a week for 30 minutes (WW) while the second half was not watered during 15 days (WD) (**Table 1)**. At the end of the two weeks of the five differential treatments, temperature was settled back at 15 °C day and night and all pots were sub-irrigated during 30 minutes once a week until the end of the experiment when plants reproduced and completed their life cycle or otherwise died.

**Table 1.**
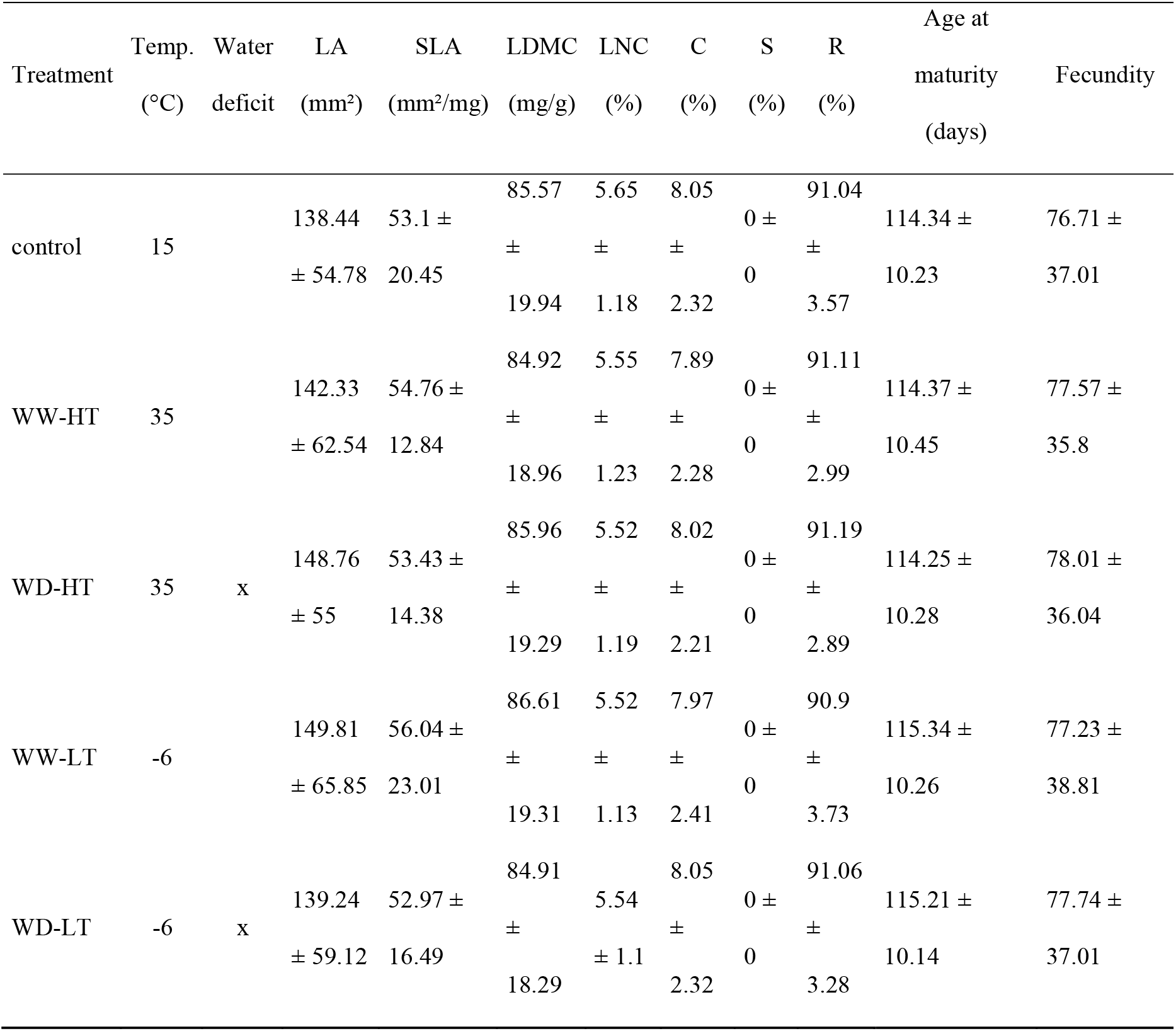
Environmental treatments and their mean effects on plant traits and CSR scores. Traits mean and standard deviation over the 30 accessions are presented for each treatment. SLA: specific leaf area; LA: leaf area; LDMC: leaf dry matter content; C: Competitive; S: Stress-tolerance; R: Ruderal. WW: well-watered; WD: water deficit; HT: hot temperature; LT: low temperature. Temp.: mean air temperature.

### Survival measurement

We estimated survival directly after the temperature treatments. An individual was considered as alive if at least the center of its rosette was still green. We estimated pre-treatment mortality by analyzing pictures of the plate the day before treatment settlement. Individuals that did not germinate or died before the treatments were discarded from the analysis.

### Leaf trait measurements and CSR scores

Two days after the end of the temperature and watering treatments, we selected 270 leaves among living individuals (2 individuals per accession and per treatment). Each leaf was rehydrated during 24 hours at 4 °C in demineralized water then weighted (Balco ME2355, France) and scanned (Epson Perfection V800, 300dpi). Then leaves were dried in an oven at 60 °C for three days and leaf dry weight determined using a balance (10^-5^g resolution, Balco ME2355, France). We measured the leaf area (LA, mm^2^) from leaf scans using ImageJ (Schneider et al., 2012). We calculated specific leaf area (SLA, mm2.g^-1^) as the ratio of leaf area to leaf dry mass. We calculate the leaf dry mass content (LDMC, mg.g^-1^) as the ratio of leaf dry mass to leaf rehydrated mass (Pérez-Harguindeguy et al., 2013). From these three leaf traits, we calculated the CSR scores from the algorithm provided by Pierce et al (2017). This algorithm implies a multivariate regression among LA, SLA and LDMC. The CSR scores obtained through this method is in accordance with scores using more traits (see Pierce et al., 2017 for more details).

### Near-infrared spectra predictions

At the end of the treatment, we acquired spectra of near-infrared reflectance of green leaves, non-destructively, using a portable spectrometer (ASD LabSpec, Malvern Panalytical, Holland, wavelength range: [780; 2500 nm]). Spectra were taken on leaves dedicated for leaf traits measurements just before harvesting and on additional individuals in order to get 12 spectra per accession and per treatment. Acquired spectra were used to predict SLA, LDMC, leaf nitrogen concentration (LNC, %), R scores and C scores for 2,160 individuals. Predictive models based on convolutional neural networks (CCNs) were developed using an independent database gathering more than 20,000 spectra and their respective reference. We evaluated the robustness of our predictions by testing their correlation with values obtained with the traditional destructive methods [**Supplementary Material S1**; **TableS2]**. Afterward, we considered predicted values superior or inferior to three median absolute values as outliers for each trait, each accession, in every treatment (Hampel, 1974). Final dataset contains traits values for 6 to 12 individuals of each of the 30 accessions in each treatment.

### Phenology and fecundity measurements

We monitored the 2,365 surviving individuals from germination to the date of the first mature and dehiscent fruit. The age at maturity was calculated as the number of days from germination to the date at which the first fruit became dehiscent. At this date, we took a picture of the inflorescence of every individual to estimate the number of fruits. We took all the pictures at the same distance from the floral stem. Based on Vasseur et al. (2018c), we first segmented the images and then shrank them in lines of crossed pixels (“skeleton”) using ImageJ (Schneider et al., 2012). Thanks to nine variables describing these skeletons and automatically measured by ImageJ, we built a linear model to estimate fecundity (n = 100, R^2^ = 0.92). This method detects aborted or non-fecundated fruits from mature and fecundated fruits (Vasseur et al., 2018c).

### Statistical analyses

We compared means of traits observed in control condition and coefficients of variation of NIRS-predicted values of traits in all treatments thanks to Tukey tests (‘Multcomp’ package, Hothorn et al., 2008). We compared cross-treatment plasticity of geographical groups through Tukey tests comparisons of the coefficient of variation across all treatments. The coefficient of variation (CV) is calculated as the total standard deviation of traits of each group across treatments divided by the cross-treatment mean. We analyzed trait plasticity in response to the treatments using linear mixed-effects models that test log-response ratios (log ratios hereafter) of traits and CSR scores as a function of geographical groups, treatments, and their interactions. Accession identity and plate identity were considered as random effects in the models. Log ratios were calculated as the logarithm of the ratio of an individual value in a given treatment and the mean value of its accession in control condition.

We analyzed the variability of performance traits (survival and number of fruits) using generalized mixed models (‘lme4’ package, Bates et al., 2015). We performed a binomial regression for survival models (logit as a link function) and a Poisson regression for fecundity models (log as a link function). We considered three fixed effects in these models: geographical origin (3 levels), treatment effect (5 levels) and their interaction. Two random effects were considered: accession identity and plate. Only one plant died in the control condition. Because most values were at the extreme of the binomial distribution in this condition, the model suffered from convergence issues. Consequently, this treatment was not compared to the others in the survival analysis.

In every model, we calculated means and standard errors of estimates with the ‘emmeans’ package (Lenth et al., 2019). We compared means between groups and between treatments with Tukey post-hoc tests (‘Multcomp’ package, Hothorn et al., 2008). We analyzed the relationship between survival and fruit production using a linear model with mean values per accessions.

## Results

### Variation of functional traits and ecological strategies across the geographical range

Under control condition, traits can be categorized into two groups: those that tend to exhibit a latitudinal gradient and those that tend to exhibit a center-margins gradient. Within the former group, geographical origin had a significant effect on age at maturity: southern accessions had a shorter lifespan than central and northern accessions (both *P* < 0.001), while northern accessions had a longer lifespan than central accessions (*P* < 0.001, **Fig. 2A**). A trend for a latitudinal gradient existed for SLA, LDMC and LNC, but no significant differences were found across geographical groups for these traits (Fig. **2BCD**). Central accessions had significantly smaller leaves than southern accessions (*P* = 0.046), and non-significantly smaller leaves than northern accessions (**Fig. 2E**). Consistent with their lower leaf area, central accessions tend to exhibit smaller C-scores and higher R-scores than southern and northern accessions, but these variations were not significant (**Fig. 2FG**). All accessions had a null S-score. Central accessions produced on average more fruits than southern and northern accessions, but the difference was only significant with northern accessions (*P* < 0.001; **Fig. 2H).**

**Figure 2.**
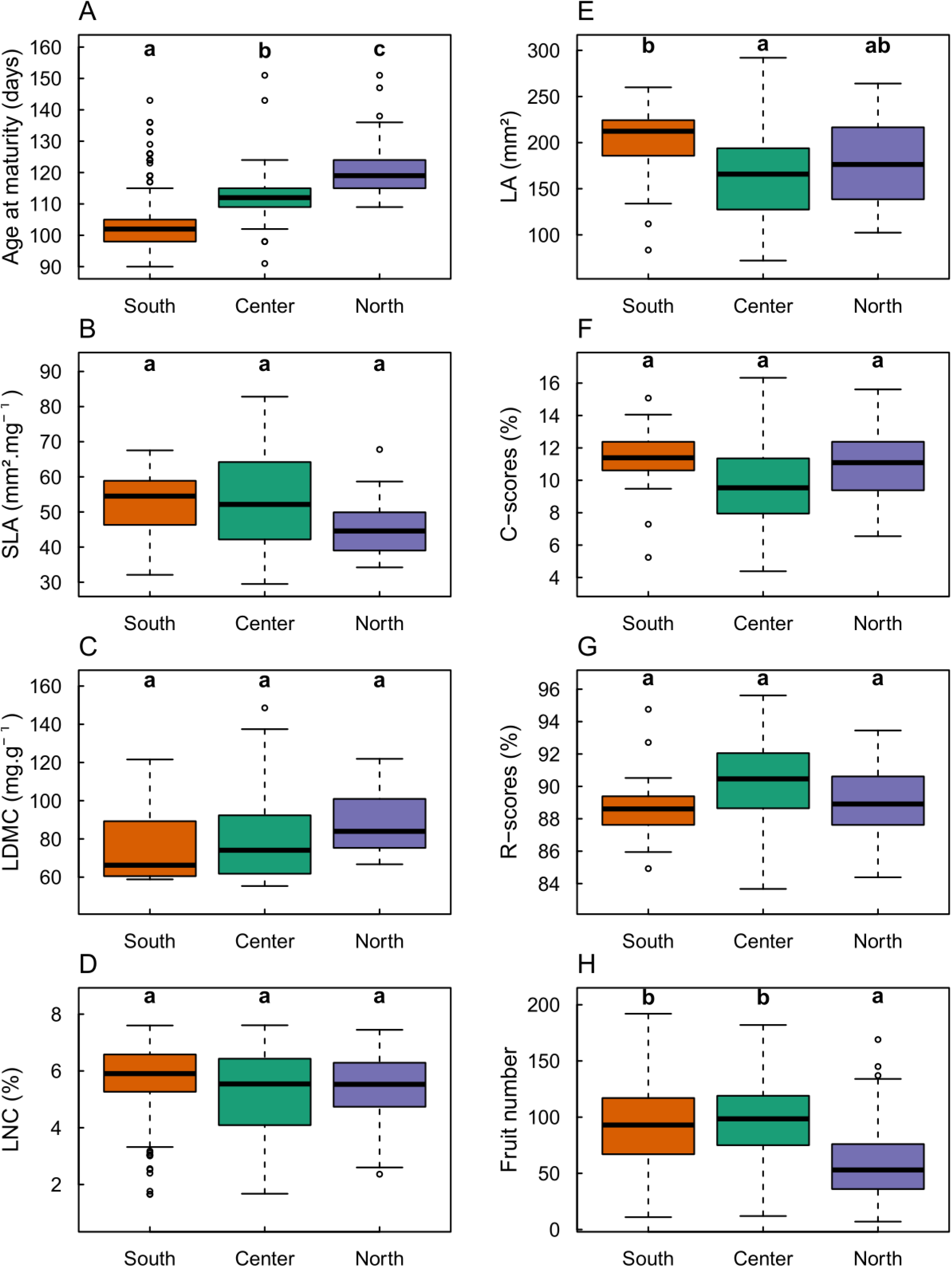
Phenotypic variation in *Control* condition across the distribution range of *A. thaliana*. Different letters indicate significant differences between geographical groups following Tukey tests at P < 0.05.

### Variation of plasticity of functional traits and ecological strategies across the geographical range

We first estimated cross-treatment trait plasticity with the coefficient of variation across five contrasted environmental conditions (control, WW-HT, WD-HT, WW-LT, WD-LT). The four traits that tended to exhibit a latitudinal gradient for trait values under control condition (age at maturity, SLA, LDMC and LNC) globally had a center-margins differentiation for trait plasticity (**Fig. 3ABCD**). For instance, central accessions had a higher plasticity of SLA across treatments than southern accessions (*P* = 0.048) and marginally higher than northern accessions (*P* = 0.054, **Fig. 3B**). Yet, the response of SLA to individual treatments exhibited more a latitudinal gradient than a center-margins gradient (**Fig. 4B**), with both southern and central accessions being more similar in their SLA log-ratio than northern accessions. Northern accessions had a higher decrease in SLA in low temperature whereas central and southern accessions had a higher increase in SLA in hot temperature conditions. Cross-treatment plasticity was not significantly different across geographical groups for age at maturity (**Fig. 3A**). Yet, the response of age at maturity to individual stress displayed a latitudinal gradient with decreasing plasticity toward the north (**Fig. 4A**). In particular, the three geographical groups differed significantly in plasticity of age at maturity in response to WD-LT. Other traits displayed more a center-margins differentiation than a latitudinal gradient. For instance, central accessions had a higher but not significant cross-treatment plasticity of LDMC (**Fig. 3C**). Central accessions had a higher coefficient of variation of LNC than northern accessions (*P* = 0.01) but did not differ significantly with southern accessions for this trait (*P* = 0.08, **Fig. 3D)**. This center-margins gradient was mainly driven by the response of LNC to WD-HT (**Fig. 4D**).

**Figure 3.**
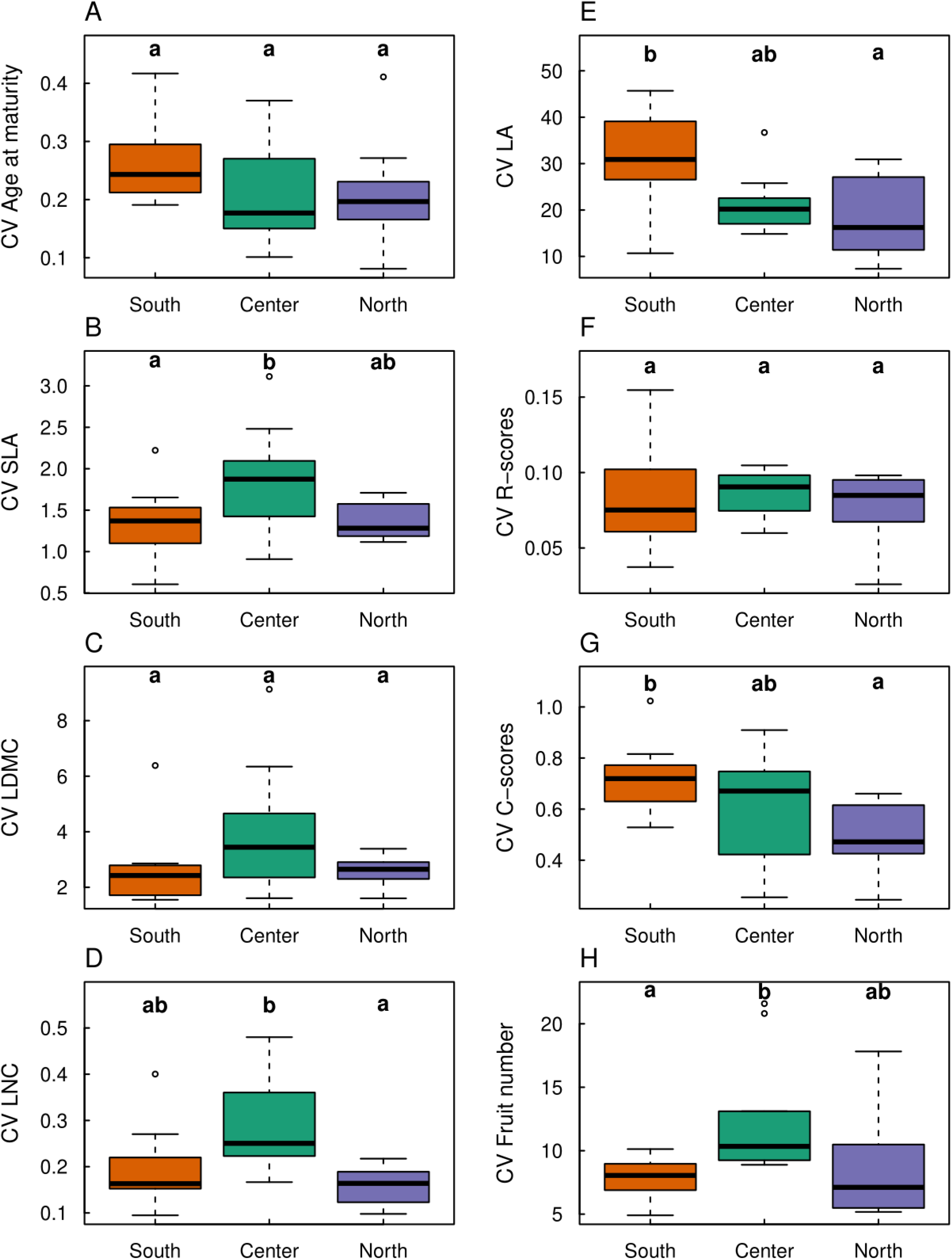
Coefficient of variation of traits across the distribution range of *A. thaliana*.Different letters indicate significant differences between geographical groups following Tukey tests at P < 0.05.

**Figure 4.**
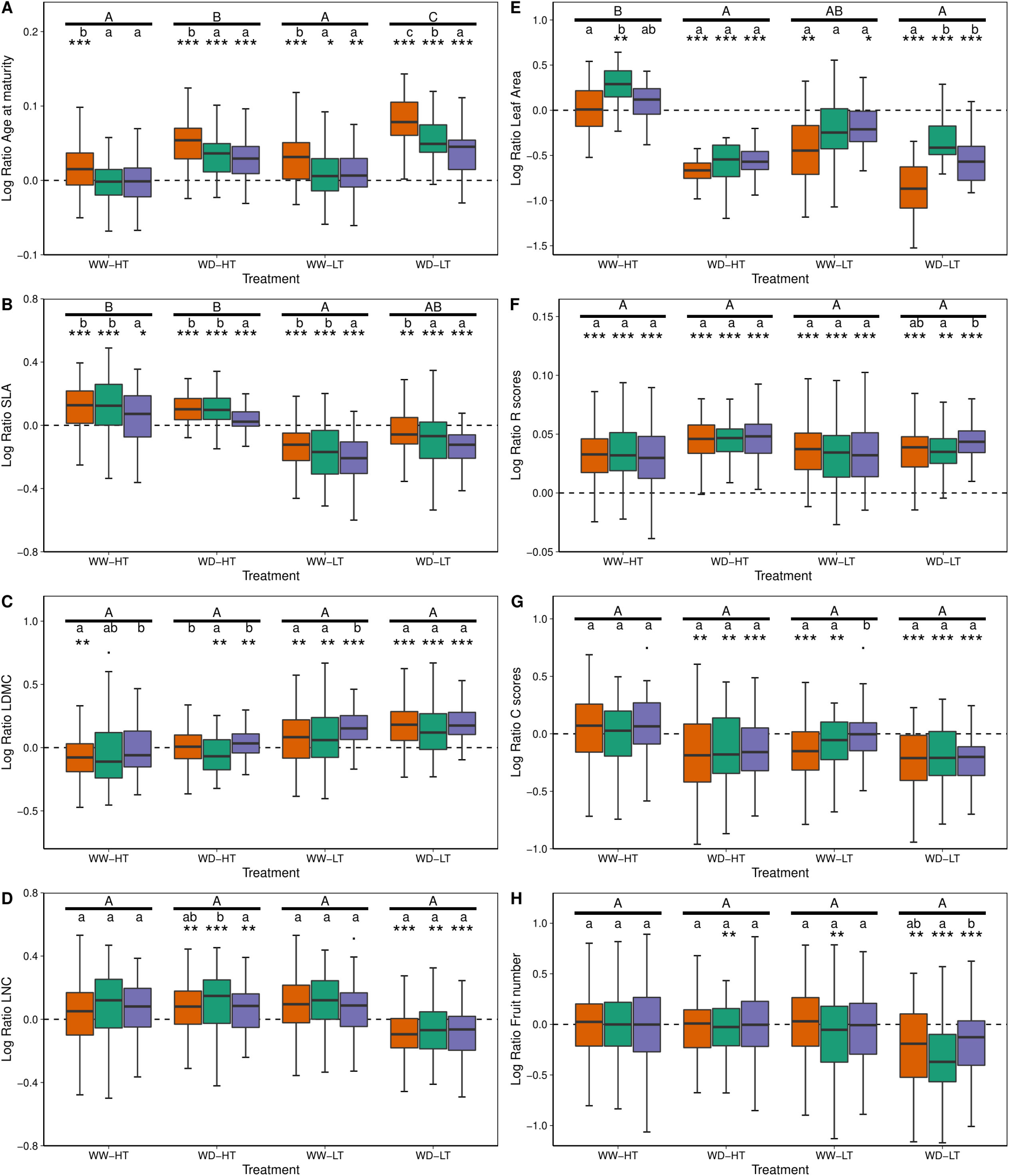
Plasticity of functional traits and CSR scores across geographical groups and treatments. Difference between mean values of Log Ratios following Tukey tests within and across treatments are indicated with lowercase letters and capital letters, respectively. Log ratios significantly different from zero following Student tests, corrected by Holm’s method, are indicated with stars (*: *P* < 0.05, **: *P* < 0.01; ***: *P* < 0.001). WW: well-watered; WD: Water deficient; HT: Hot temperature; LT: Low temperature.

Among the four traits that tended to exhibit a center-margins gradient in non-stressing conditions (LA, C and R scores, and fruit number), only fruit number also had a center-margins gradient for trait plasticity (**Fig. 3H**). Central accessions had a significantly higher coefficient of variation of fruit number than southern accessions (*P* = 0.04) and slightly higher than northern accession even if not significantly different (*P* = 0.15). Central accessions are the only accessions to produce less fruits in WD-HT and in WW-LT conditions than in control. (**Fig. 4H**). In contrast to fruit number, LA and C-scores exhibited a significant latitudinal gradient for cross-treatment plasticity. Northern accessions had a significantly lower plasticity of LA than southern accessions (*P* = 0.009), central accessions having an intermediate but not significantly different coefficient of variation on this trait (**Fig. 3E**). Yet, this cross-treatment plasticity pattern hides contrasted responses to individual stresses: plasticity of LA displayed a clear center-margins gradient only in WW-HT and WD-LT, but it exhibited a latitudinal gradient in response to WD-HT and WW-LT (**Fig. 4E**). Similar to LA, C-scores had a smaller coefficient of variation in northern than in southern accessions across treatments (*P* = 0.006), central accessions having an intermediate but not significantly different coefficient of variation for this trait (**Fig. 3G)**. Yet, only northern accessions exhibited a significantly different response of C-scores to WW-LT when looking at individual treatment effect (**Fig. 4G**). The cross-treatment plasticity of R-scores exhibited no differences between geographical groups (**Fig. 3F**), although it exhibited significantly different response of northern accessions to WD-LT (**Fig. 4F**). All accessions had a null S-score in every treatment.

### Geographical origin effects on survival

Survival of accessions varied significantly among treatments (*P* < 0.001). In particular, WW-LT (79.9% of survival), WD-HT (72.2% of survival) and WD-LT (52.4% of survival) were associated with a significantly weaker probability of survival than WW-HT (99.2% of survival) (*P* = 0.02; *P* = 0.0003; and *P* < 0.001 respectively). A single individual died in Control (99.8% of survival), likely unrelated to adaptation to such conditions. Survival of accessions varied significantly across geographical groups (*P* < 0.001), which globally exhibited a latitudinal gradient. Among all treatments, northern accessions survived significantly more than central (*P* = 0.01) and southern accessions (*P* = 0.001), consistent with the significant interaction between geographical group and treatment (*P* < 0.001). In WW-LT, central accessions survived more than southern accessions (*P* = 0.007). Northern accessions survived significantly more than the central and southern accessions in the cold treatments (WW-LT and WD-LT, resp. *P* < 0.001). Moreover, northern accessions survived significantly more than central accessions (*P* = 0.046) in WD-HT, but they were not different from southern accessions (*P* = 0.06) (**Fig. 5A**).

**Figure 5.**
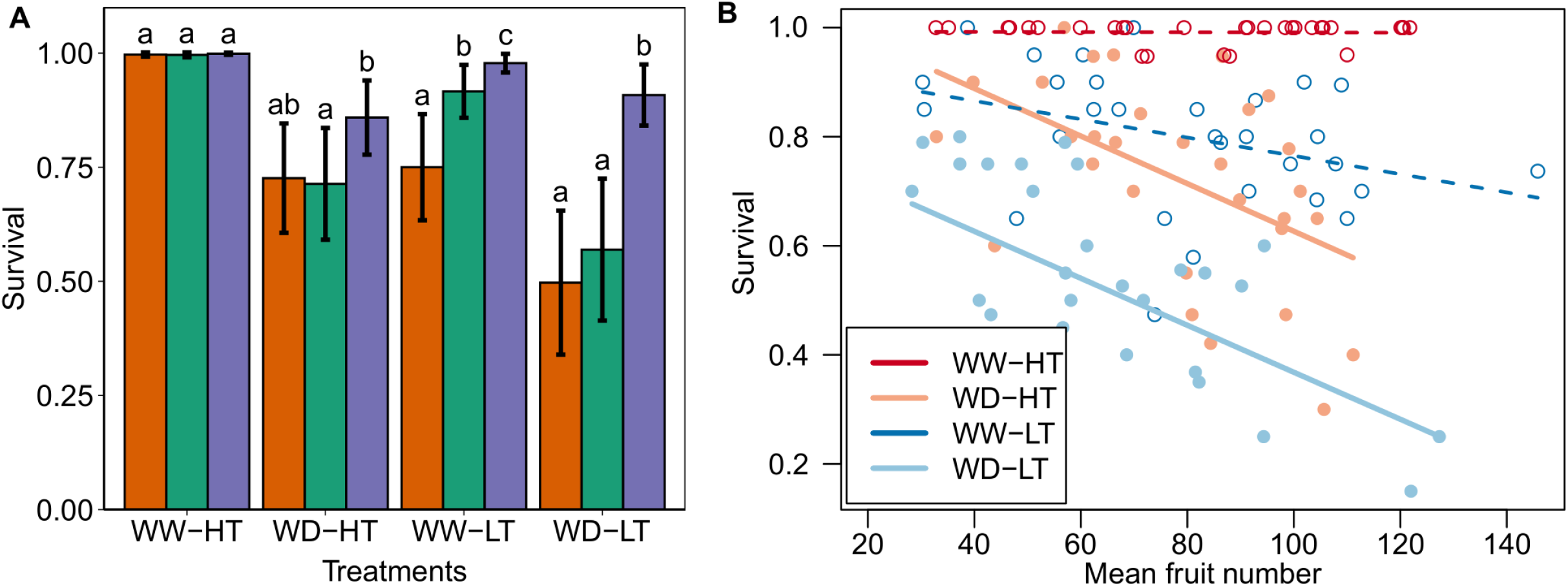
A) Survival rate among treatments. Different letters indicate significant differences between geographical groups following Tukey tests at P < 0.0.05 within each treatment. B) Relationship between survival of accessions (n = 30) and fruit production across treatments. Linear regression lines are indicated. Dashed lines and empty points indicate a slope not significantly different from zero. The relationship under control condition was not significant and is not shown. WW: well-watered; WD: water deficit; HT: hot temperature; LT: low temperature.

Survival was significantly negatively related to fruit number under WD-HT and WD-LT (R^2^ = 0.75; *P* < 0.001) but the slope of the relationship was not significantly different from zero under control, WW-HT and WW-LT. In other words, under WD conditions, accessions with low fecundity survived more than accessions with high fecundity (**Fig. 5B, [Supplementary Information Fig. S2**).

## Discussion

This study dissects functional variation at the intraspecific level within different environments and across the distribution range of a widespread species. We expected two main types of geographic mean trait variation patterns across *A. thaliana* distribution; either a latitudinal gradient or a differentiation between the center and the margins of the distribution. The studied traits and their plasticity were correlated but they exhibited various patterns of geographic variation. We discuss the consequences for variation in individual performance and local plant adaptation across the range.

In Europe, *Arabidopsis thaliana* faces very contrasted climates (Hoffmann, 2002), which are expected to constitute strong yet variable natural selection pressures throughout its distribution range (Kaplan et al., 2004; Swindell et al., 2007; Vile et al., 2012). Coherently, part of our results supports a latitudinal gradient in functional variation across the distribution range of *A. thaliana*. In non-stressful conditions for plant growth, age at maturity, specific leaf area, leaf dry matter content, and leaf nitrogen concentration vary along this latitudinal gradient. These traits are closely associated with the leaf economics spectrum (Wright et al., 2004). Our results support a latitudinal gradient in resource-use strategies, from acquisitive resource-use strategy for southern accessions (characterized by short lifespan, thin leaves with high LNC and photosynthetic rate) to conservative resource-use strategy in northern accessions (characterized by long lifespan, thick leaves with low LNC and photosynthetic rate). An abundant literature in *A. thaliana* supports this functional gradient associated with latitude in Europe (Stenøien et al., 2002; Stinchcombe et al., 2004; Hopkins et al., 2008; Vasseur et al., 2012; Debieu et al., 2013; Vasseur et al., 2018a; Exposito-Alonso, 2020). By contrast, Sartori et al., (2019) found that both southern and northern accessions displayed a conservative resource-use strategy. Here, we show that northern accessions had a higher survival rate at low temperature than central and southern accessions. This suggests that conservative resource-use strategies selected in cold climates in northern areas of Europe is associated with an optimization of survival to freeze. Surprisingly though, northern accessions had also a higher survival rate than southern and central accessions in the hot temperature treatment. We can hypothesize that the metabolic pathways associated to a better survival in dehydration caused by freeze could also be efficient for a better survival under water deficit conditions (Sanada et al., 2007; Suprasanna et al., 2016; Gillespie and Volaire, 2017; Bristiel et al., 2018). However, the inverse is not true, southern accessions being the most vulnerable to nocturnal freezing. A possible explanation is that stress escaping (Ludlow, 1989) is closely associated with acquisitive resource-use strategies that are selected in the southern area of the distribution range of *A. thaliana*. However, in southern populations, the two opposite strategies are coexisting with four southern accessions surviving more than the other southern accessions. Interestingly, these four accessions are from the same genetic lineage: the relict group ([**Supplementary Information Table S1**], 1001 Genomes Consortium, 2016). This ancient genetic lineage is associated with stress-tolerance in Spain (Lee et al., 2017). Modern Spanish accessions present a short life cycle following spring germination (our results; Assmann, 2013; Exposito-Alonso, 2020) that is strongly associated with an acquisitive resource-use strategy. Oppositely, northern accessions from Scandinavia where low temperatures and short spring season do not allow for a rapid life cycle strategy, plants are selected for high tolerance strategy to resist winter conditions (Bartlett et al., 2014; Delzon, 2015; Exposito-Alonso, 2020). Underpinning this tolerance/avoidance trade-off, southern survivors under stressful conditions increased more their life duration than central and northern accessions. This corroborates the study of Exposito-Alonso et al. (2020) who showed that Spanish accessions had a more plastic life cycle than Scandinavian strict winter cycler accessions. These results also corroborate the genetic correlation between water use efficiency and life span found in previous studies (Mckay et al., 2003): increasing water use efficiency through phenotypic plasticity may constrain the life cycle of individuals to be longer.

The center-margins gradient of abiotic stress hypothesis, which posits that less suitable environments occur at the peripheries (Holt, 2009), has been discussed on numerous species (Sexton et al., 2009; Pironon et al., 2017). In *A. thaliana*’s distribution range, mean annual precipitation exhibits a bell-shaped curve with latitude. Northern and southern populations encountering less precipitations than central populations **[Supplementary Information Fig. S1]**. Low precipitations are expected to reduce the variance of phenotypes associated with water-stress resistance and may limit phenotypic plasticity (Valladares et al., 2007; Palacio-López et al., 2015; Stotz et al., 2021). In parallel, northern and southern parts of the distribution range of *A. thaliana* encounter more seasonal variation of temperature and precipitation **[Supplementary Information Fig. S1].** Fluctuating conditions, when predictable, are expected to select for plastic phenotypes (Lázaro-Nogal et al., 2015; Leung et al., 2020; Stotz et al., 2021). Our results show a general trend for a weaker cross-treatment plasticity in peripheral accessions than central accessions. This could be explained by the cost of phenotypic plasticity, being higher in stressful conditions associated with fewer precipitations (van Kleunen and Fischer, 2005; Molina-Montenegro and Naya, 2012; Nicotra et al., 2015). A weaker or an absence of phenotypic plasticity as well as a stronger genetic determinism independent of the climate at ecological margins may be adaptive (Ghalambor et al., 2007; Murren et al., 2015; Palacio-López et al., 2015; Acasuso-Rivero et al., 2019; Pfennig, 2021). Indeed, only the central populations showed a significant fecundity reduction in stress conditions. This result is also in accordance with Exposito-Alonso et al. (2018) who showed that similar alleles involved in drought resistance are under selection at both latitudinal margins of Europe in *A. thaliana*. Our results demonstrate stress-tolerance in *A. thaliana* whereas all of the accessions had a null S-score in our estimations, based on Pierce et al.’s methodology (2017). This result questions the use of such classification at the intraspecific level. We suggest improving the CSR classification by using phenotypic traits and performances actually measured under competition, disturbance and stress conditions.

The reduction of plasticity at both margins of the distribution range may also be associated with the evolutionary history of the species. Actual marginal populations of *A. thaliana* derived from the European colonization of a genetic lineage from central Europe and its admixture with northern and southern populations (Lee et al., 2017). The colonization of margins may have been accompanied with both high cumulative foundation effects and directional selection, limiting thus the phenotypic variability at both opposite margins (Kirkpatrick and Barton, 1997; Sagarin and Gaines, 2002; Bridle and Vines, 2007; Eckert et al., 2008; Sexton et al., 2009; Luo et al., 2015; Pironon et al., 2017; Hämälä et al., 2018). Accordingly, leaf area, C-scores, R-scores and fruit number are more similar among the two opposite margins than with the central accessions. Our results suggest that trait values associated with colonization (number of fruits and R-scores) are higher in central accessions than peripheral ones. Moreover, traits related to competitive ability (leaf area and C-scores) are lower in central accessions than peripheral ones. This result is quite counter-intuitive regarding theoretical expectations that abundance in central populations should be higher than in marginal populations, selecting thus for a higher competition ability at the center than at the margins of a distribution range (Brown, 1984; Brown et al., 1995; Holt, 2009; Sexton et al., 2009; but see Pironon et al., 2017). We show that central accessions invest more in fecundity than peripheral accessions who invest more in resistance to stress. Grounded on ecological theories regarding the existence of a colonization/competition trade-off (Levins and Culver, 1971; Hastings, 1980; Turnbull et al., 1999; Yu and Wilson, 2001; Cadotte et al., 2006), we can thus hypothesize that central accessions exhibit traits that optimize seed dispersal and colonization, perhaps also at the expense of competitive or stress-coping ability. Indeed, we confirm the classical trade-off described at the interspecific level between survival to stress and fecundity (Muller-Landau, 2010; D’Andrea et al., 2013). It may be thus interesting to experimentally test this competition/colonization ability differentiation across the distribution range in order to better understand how phenotypes evolved through the evolutionary history of *A. thaliana* (e.g. Lorts and Lasky, 2020).

All in all, our work confirmed hypothesized trends on how ecological strategies vary across the geographic distribution of a species. More importantly, we show that both climate variations and evolutionary history shaped the actual phenotypic diversity in this model species, leading to a latitudinal and a center-margins differentiation respectively depending on the nature of the traits. The latitudinal gradient was associated with an acquisition/conservation trade-off, tightly linked to a temperature gradient along european latitudes. At the opposite, the center-margins differentiation was more associated with a competition/colonization trade-off potentially due to the demographic history of this species. Our findings thus point out the importance of considering the structured phenotypic variability of species to understand the ecology and evolution of species’ ranges rather than comparing species using their mean trait values only. This is particularly important to better predict distribution range future evolution.

## Acknowledgements

We are very grateful to Justine Floret and Amelie Emmanuel for their help on measurements. We also thank the technical platform “Plateforme des Terrains d’Expérience du LabEx CeMEB,” (CEFE, CNRS) in Montpellier and particularly Pauline Durbin and Thierry Matthieu for their precious advices. This work was supported by the European Research Council (ERC) Starting Grant Project “Ecophysiological and biophysical constraints on domestication in crop plants” (Grant ERC-StG-2014-639706-CONSTRAINTS).

## Supplementary Information

**Table S1**. List of accessions and their geographical coordinates. **Figure S1.** Description of climate of origin of the three geographical groups compared to the diversity of climate from the complete set of 1135 accessions available in the 1001genome project. Data were downloaded from https://chelsa-climate.org/ **Supplementary Material S2.** Predictive models’ development. **Table S2.** Performances of models predicting leaf traits from near infrared spectroscopy predictive models for in-sample and cross-validation sets of (N: number of samples, RMSE: Root Mean Square Error, R2: Coefficient of Determination). **Figure S2.** Relationship between survival of accessions among geographical groups in WD-HT and WD-LT. Labels referred to accessions ID (see Table S1).

## Notes

### Competing Interest Statement

The authors have declared no competing interest.

