## Supplementary material for "Into the range: a latitudinal gradient or a center-margins differentiation of ecological strategies in *Arabidopsis thaliana*?": Table S1

**Table S1.** Set of accessions used in this study. Accessions ID and names came from 1001genome project denomination.

| Group | Accession ID | Name | Latitude | Longitude | Altitude (m) | Genetic group |
|---|---|---|---|---|---|---|
| Centre | 6898 | An-1 | 51.21 | 4.4 | 5 | admixed |
| | 7213 | Ler-0 | 47.9 | 10.87 | 630 | admixed |
| | 7382 | Utrecht | 52.09 | 5.11 | 4 | admixed |
| | 7298 | Pi-0 | 47.04 | 10.51 | 2395 | central europe |
| | 8311 | In-0 | 47.5 | 11.5 | 1476 | central europe |
| | 6915 | Ei-2 | 50.3 | 6.3 | 513 | germany |
| | 6897 | Ag-0 | 45.0 | 1.3 | NA | western europe |
| | 6958 | Ra-0 | 46.0 | 3.3 | 316 | western europe |
| | 6959 | Rennes-1 | 48.5 | -1.41 | 89 | western europe |
| | 7092 | Com-1 | 49.41 | 2.823 | 45 | western europe |
| Nord | 6009 | Eden-1 | 62.8 | 18.17 | 90 | north sweden |
| | 6177 | TAL 03 | 62.63 | 17.69 | 137 | north sweden |
| | 6184 | TBA– 01 | 62.88 | 18.45 | 21 | north sweden |
| | 6209 | TEDEN 02 | 62.88 | 18.18 | 90 | north sweden |
| | 6244 | TRA01 | 62.91 | 18.47 | 65 | north sweden |
| | 8376 | Sanna-2 | 62.69 | 18.0 | 78 | north sweden |
| | 6074 | A–r-1 | 56.45 | 16.13 | 2 | south sweden |
| | 8240 | Kulturen-1 | 55.70 | 13.19 | 26 | south sweden |
| | 9057 | VinslAv | 56.1 | 13.91 | 22 | south sweden |
| | 9470 | Tur-4 | 57.65 | 14.80 | 273 | south sweden |
| Sud | 6911 | Cvi-0 | 15.11 | -23.61 | 270 | relict |
| | 9549 | IP-Hum-2 | 42.23 | -3.69 | 903 | relict |
| | 9600 | IP-Vis-0 | 39.85 | -6.04 | 338 | relict |
| | 9947 | Ped-0 | 40.74 | -3.9 | 1007 | relict |
| | 6970 | Ts-1 | 41.71 | 2.93 | 116 | spain |
| | 8357 | Pla-0 | 41.5 | 2.25 | 145 | spain |
| | 9507 | IP-Coa-0 | 38.45 | -7.5 | 190 | spain |
| | 9537 | IP-Cum-1 | 38.07 | -6.66 | 596 | spain |
| | 9544 | IP-Gua-1 | 39.4 | -5.33 | 694 | spain |
| | 9943 | Cdm-0 | 39.73 | -5.74 | 444 | spain |
