## Supplementary material for "Into the range: a latitudinal gradient or a center-margins differentiation of ecological strategies in *Arabidopsis thaliana*?": Table S2

**Table S2:** Performances of models predicting leaf traits from near infrared spectroscopy predictive models for in-sample and cross-validation sets of (N: number of samples, RMSE: Root Mean Square Error, $R^2$: Coefficient of Determination).

| Trait | Internal validation | | | External validation | | |
|---|---|---|---|---|---|---|
| | N | RMSE | $R^2$ | N | RMSE | $R^2$ |
| SLA | 1039 | 7.667 | 0.842 | 294 | 6.67 | 0.51 |
| LDMC | 1027 | 16.099 | 0.860 | 294 | 0.99 | 0.68 |
| LNC | 460 | 0.533 | 0.925 | NA | NA | NA |
| R-scores | 1015 | 4.656 | 0.879 | 294 | 2.48 | 0.29 |
| C-scores | 1015 | 2.955 | 0.903 | 294 | 1.88 | 0.11 |
