## Supplementary figures and images for "Into the range: a latitudinal gradient or a center-margins differentiation of ecological strategies in *Arabidopsis thaliana*?"

### Figure S1

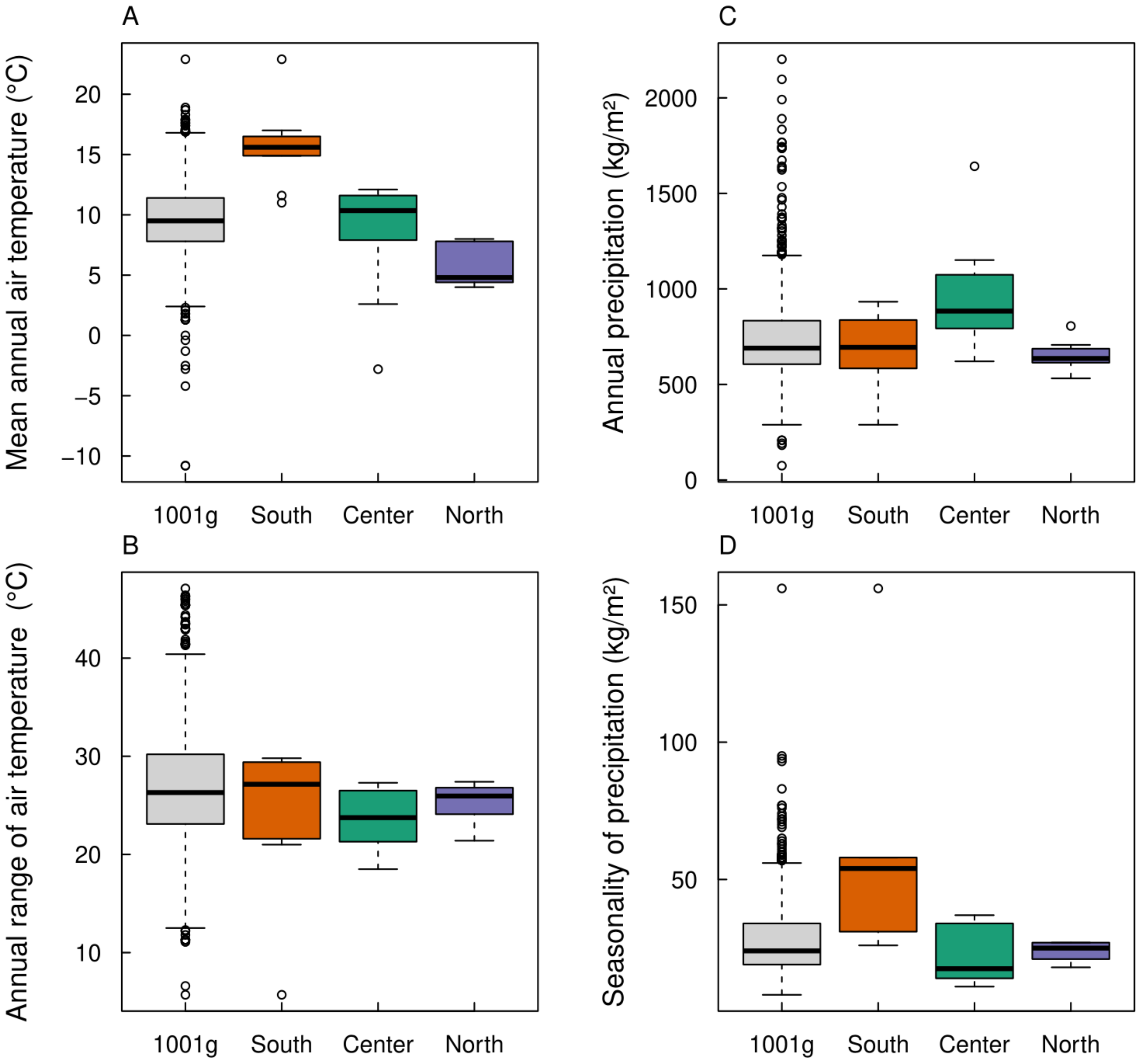

### Figure S2

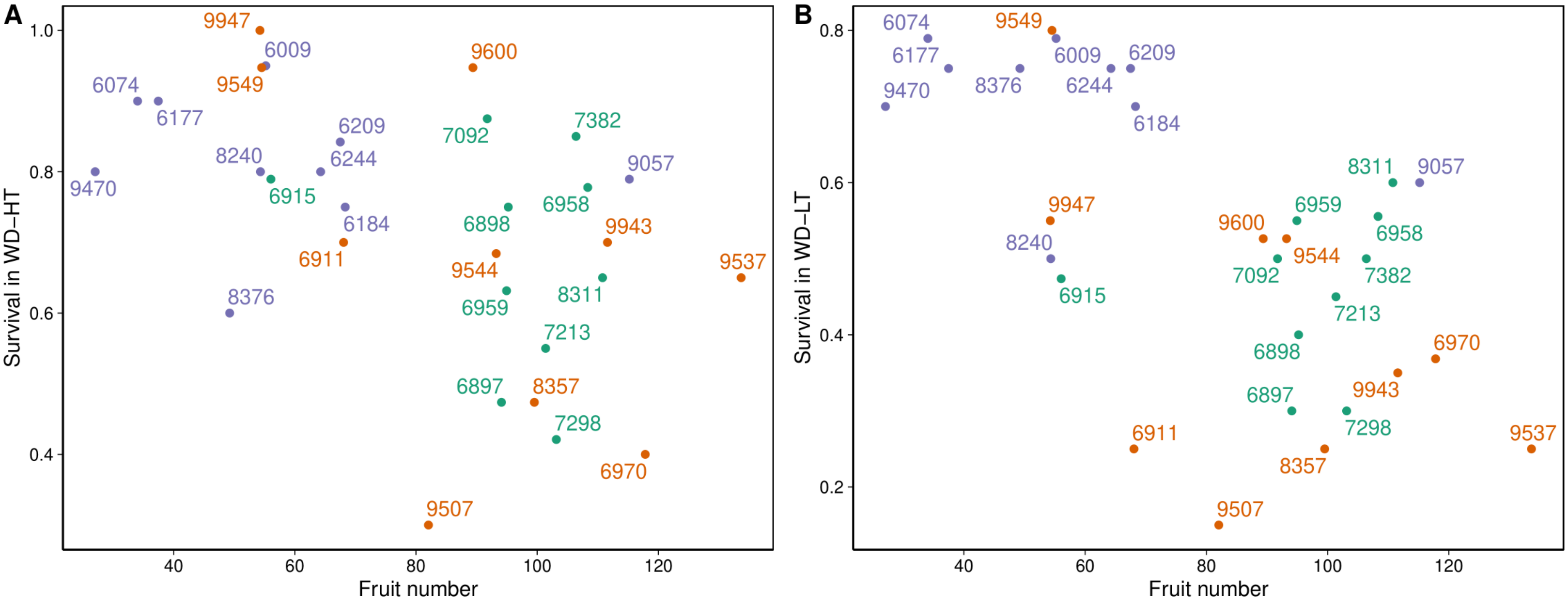
