## Supplementary Material S2 for "Into the range: a latitudinal gradient or a center-margins differentiation of ecological strategies in *Arabidopsis thaliana*?"

**Supplementary Material S2.** Predictive models' development

For the calibration of these models, an independent database gathering more than 20000 spectra and their respective reference values was used. Pretreatments, calibration and validation were carried out using python language (v3.6, https://www.python.org) with a Keras framework (v2.1.5, https://keras.io/) and a TensorFlow backend (v1.6.0, https://www.tensorflow.org/). Samples were divided into a calibration set (75%) and an internal validation set (25%) using the Kennard-Stone algorithm.

First, we took a data (i.e. sample) augmentation approach to double the size of the calibration set and improve generality of the network's learning. For each original sample, 30 synthetic spectra were generated using a combination of random translation and rotation of the original spectra. Then spectra presenting absorbance values higher than 1 or lower than 0 were discarded. This allowed to expand the original 61 spectra to more than 800 spectra. This data augmentation with noise addition is a common technique used in deep learning to reduce overfitting of small datasets.

Secondly, we conducted a feature data (i.e. spectral) augmentation on the calibration and validation sets. We applied an all-possibilities approach, combining and keeping the different pretreated spectra, including the original one. For each sample, feature augmentation was applied by generating 12 new spectra using pretreatments based on Haar transform, Gaussian derivatives, SVG, SNV, and different degrees of MSC.

A convolutional neural network composed of three convolutional layers followed by two dense layers was fitted to the calibration data. Mean squared error was used as the loss function. In order to avoid overfitting, two batch normalizations and a dropout of 15% of features were applied respectively between layers. The model was calibrated using a three-fold cross validation approach. Model performance was evaluated using Root Mean Square Error (RMSE) and Coefficient of Determination (R2) (**Table S2)**.
